# Nanopore Direct RNA Sequencing Enables Reproducible, Site-Resolved Pseudouridine Quantification in Human Ribosomal RNA

**DOI:** 10.64898/2026.06.08.730855

**Authors:** Baudouin S de Préval, Laurence Faucher-Giguère, Maxime Duval, Virginie Marchand, Pavan Lakshmi Narasimha, Nehal Thakor, Yuri Motorin, Sherif Abou Elela, Michelle S Scott

## Abstract

Pseudouridine is the most abundant post-transcriptional modification in human ribosomal RNA, with over 110 annotated sites and variable stoichiometry across biological contexts. Existing quantification methods are low-throughput or constrained to predefined panels. We benchmarked nanopore direct RNA sequencing using the Dorado v5.1 model against mass spectrometry–validated sites in human liver tissue, induced pluripotent stem cells, and HeLa cells. Nanopore sequencing detected 95 of 117 validated sites and accurately quantified stoichiometry at 85% of sites with high reproducibility. Low GC-content environments were the primary source of failure. These results establish nanopore sequencing as a scalable tool for epitranscriptomic pseudouridine profiling.

## Introduction

From its initial description as a “fifth” nucleotide in 1951 [1] to its established roles in RNA structure and function, pseudouridine has been studied for decades as a conserved post-transcriptional RNA modification [2–4]. This C5-glycoside isomer of uridine occurs in multiple RNA classes, including transfer RNA, messenger RNA, and ribosomal RNA, where it contributes to RNA stability, folding, and molecular interactions within the ribosome [5,6]. In ribosomal RNA (rRNA), pseudouridine is one of the two most abundant chemical modifications, together with 2′-O-methylation, with more than 110 sites currently annotated in human ribosomes [7,8]. Most rRNA pseudouridine positions are constitutively modified; however, a subset exhibits fractional occupancy that can vary across biological contexts. Alongside rRNA sequence variation and ribosomal protein heterogeneity, variation in rRNA modification stoichiometry constitute a triptych that has been proposed to contribute to ribosome heterogeneity [9–14]. Although loss of individual pseudouridines is often tolerated, altered stoichiometry at specific sites can affect ribosome composition and translational fidelity and has been associated with selective translational programs in cancer and other disease contexts [15–18]. For example, hypo-pseudouridylation of 18S rRNA at position U1248 has been reported in a substantial fraction of colorectal cancers and linked to increased ribosome biogenesis and tumor cell invasiveness [19]. More broadly, changes in rRNA modification stoichiometry have been reported across multiple cancer types and have also been observed to vary between tissues and physiological conditions, highlighting how modulation of pseudouridine levels may influence local ribosomal structure and interactions with specific messenger RNAs [20].

Despite this accumulating evidence, the functional consequences of rRNA pseudouridine heterogeneity continue to be poorly understood, primarily due to the difficulty in accurately measuring modification levels at scale. Quantitative characterization of pseudouridine occupancy has largely remained qualitative or restricted to a limited number of predefined sites, underscoring the need for a comprehensive evaluation of modification stoichiometry across the transcriptome. Early strategies to address this gap generally relied on the selective reactivity of N-cyclohexyl-N′-β-(4-methylmorpholinium)ethylcarbodiimide (CMC) with pseudouridine [21,22]. Under alkaline conditions, CMC forms an adduct with pseudouridine that stalls reverse transcription, generating characteristic termination signatures during cDNA synthesis. While these signatures enable sensitive site identification, they provide only indirect and often limited quantitative information on modification stoichiometry.

To obtain numerical estimates of pseudouridine occupancy, mass spectrometry–based approaches were adapted to RNA through cyanoethylation chemistry, allowing pseudouridine to be distinguished from uridine that otherwise share an identical atomic mass [23,24]. Although this strategy provides accurate stoichiometric measurements, it is labor-intensive and difficult to scale across samples or conditions. Advances in high-throughput sequencing subsequently led to CMC-derived methods such as Ψ-seq [25] and related approaches [26–29], culminating in HydraPsiSeq [30], which exploits the resistance of pseudouridine to hydrazine/aniline cleavage. In this method, cleavage at unmodified uridines generates characteristic 5′ read ends, while pseudouridine positions show reduced cleavage and lower local read counts, enabling calculation of a quantitative PsiScore. However, HydraPsiSeq relies on targeted amplification of predefined rRNA regions prior to sequencing, which constrains analysis to known sites, introduces PCR-associated biases, and precludes de novo discovery of pseudouridine positions—limitations that become particularly relevant in densely modified RNAs such as rRNA.

In parallel to chemical and mass spectrometry–based approaches, direct RNA sequencing by nanopore technology has recently emerged as an alternative strategy for RNA modification analysis. This approach infers nucleotide identity and modification status from characteristic disruptions in the ionic current generated as individual RNA molecules translocate through a nanopore embedded in a polarized membrane. For epitranscriptomic applications, modification signatures can be detected either indirectly through systematic basecalling errors, such as uridine-to-cytidine mismatches associated with pseudouridine [31], or by exploiting additional signal features, including current intensity, dwell time, and signal variance, using supervised or semi-supervised machine learning models [32–34]. Oxford Nanopore Technologies (ONT) has further incorporated modification-aware basecalling frameworks that enable direct prediction of RNA modifications from raw sequencing data [35,36]. These developments offer the prospect of single-molecule, site-resolved quantification of RNA modifications and the simultaneous assessment of multiple modification types on native RNA strands. However, their application to densely modified and highly structured RNAs remains challenging. Recent benchmarking studies have reported elevated false-positive rates and context-dependent inaccuracies, particularly at sites embedded in complex secondary structures or proximal to other modifications. Of note, these studies have focused primarily on messenger RNA or synthetic constructs, leaving rRNA, the most densely modified, highly structured, and repetitive cellular RNA, largely uncharacterized in this context. [37,38]. Systematic evaluation of nanopore-based pseudouridine detection in native human rRNA against orthogonally validated stoichiometric ground truth across biologically distinct sample types therefore remains lacking.

Here, we assess the capacity of current ONT direct RNA sequencing technology to detect and quantify pseudouridylation in human ribosomal RNA. Using the pseudouridine-aware Dorado basecalling model (v5.1), we sequenced rRNA from human liver tissue, human induced pluripotent stem cells, and HeLa cells and benchmarked nanopore-derived estimates against orthogonal measurements from mass spectrometry and HydraPsiSeq (Figure 1A). Focusing on a curated set of mass spectrometry–validated pseudouridine sites, we quantified detection rates, reproducibility across technical replicates and batches, and the sequence/structural contexts associated with agreement or disagreement between methods.

**Figure 1.**
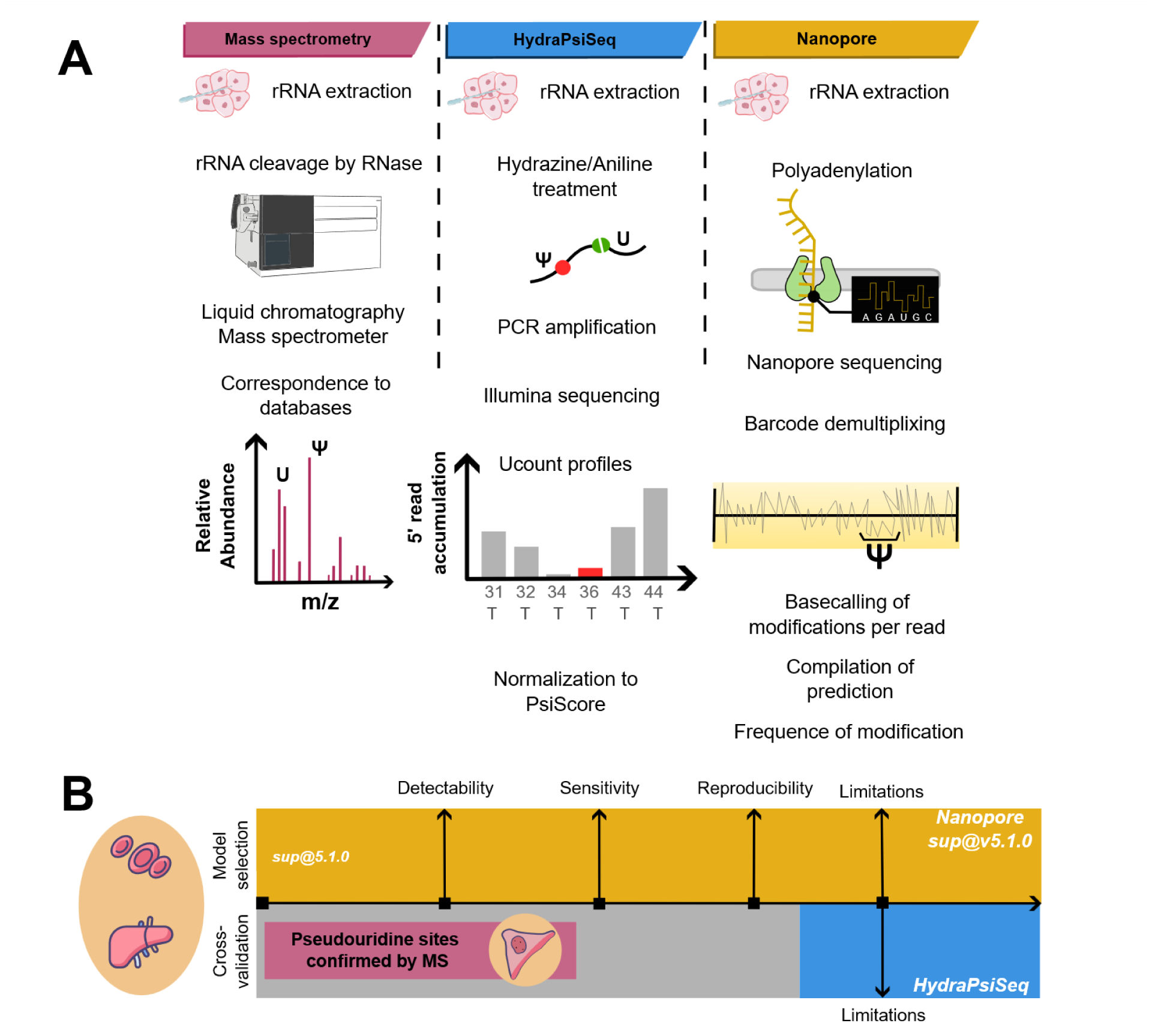
Comparison of experimental pipelines and assessment framework for rRNA pseudouridine (Ψ) detection. **(A)** Schematic overview of the three complementary approaches used to identify and quantify pseudouridine in ribosomal RNA (rRNA). In the mass spectrometry workflow, rRNA is extracted, enzymatically cleaved, and analyzed by liquid chromatography–mass spectrometry to detect modified and unmodified nucleosides based on their mass-to-charge signatures. In HydraPsiSeq, rRNA undergoes selective hydrazine/aniline cleavage followed by reverse transcription, PCR amplification and Illumina sequencing; cleavage-dependent 5′ read-accumulation profiles are used to calculate normalized PsiScores that estimate modification frequencies. In nanopore sequencing, native rRNA molecules are sequenced directly after a polyadenylation step, barcode demultiplexing enables sample-specific signal extraction, and model-based basecalling generates per-read Ψ predictions that are aggregated into site-specific modification frequencies. **(B)** Benchmarking framework used to assess performance across methods. Biological samples analyzed in this study are shown on the left. At the top, the nanopore basecalling model applied (<u>sup@v5.1.0</u>) and the evaluation criteria (detectability, sensitivity, reproducibility, and method-specific limitations) are indicated. Previously published mass-spectrometry-validated Ψ sites in HeLa rRNA served as positive controls for cross-validation. Nanopore-derived modification frequencies were compared with matched HydraPsiSeq measurements to evaluate concordance and define strengths and limitations of each method.

## Results

### Definition of a mass spectrometry–validated rRNA pseudouridine reference and nanopore detection framework

To define the set of rRNA positions evaluated by nanopore sequencing, we compiled a reference dataset of pseudouridine sites supported by orthogonal measurements. Mass spectrometry–validated positions reported in two independent studies were integrated to generate a non-redundant repertoire of rRNA pseudouridine sites with unambiguous experimental support. This reference dataset comprised 117 positions distributed across human rRNA (62 in 28S, 53 in 18S, and 2 in 5.8S) and was used throughout the study to focus nanopore-based analyses on the quantification of robustly validated sites. Using this reference framework, we generated direct RNA nanopore sequencing datasets from rRNA isolated from human liver tissue and human induced pluripotent stem cells (hIPSCs), providing two distinct biological contexts for method evaluation (Figure 1B). To enable multiplexed analysis and assessment of batch-level reproducibility, samples were barcoded [39] prior to sequencing and processed as independent technical replicates. Sequencing yielded at least 20,000 reads per sample except for one hIPSC sample that yielded 17,874 reads, still sufficient for long-read epitranscriptomic analyses [40,41]. Reads were aligned to reference human rRNA sequences, including annotated 28S and 18S variants, and pseudouridine calls were derived from raw signal using the modification-aware Dorado basecaller (rna004_130bps_sup@5.1).

We first examined pseudouridylation frequencies reported by ONT at the 117 rRNA sites validated by mass spectrometry. Across all predicted positions, the distribution of estimated modification frequencies showed a progressive depletion of non–MS-validated sites as frequency increased, with a clear separation emerging above approximately 15% (Figure 2A). Applying a minimum frequency threshold of 15% recovered the majority (89%) of MS-validated pseudouridine sites (green density), while substantially reducing low-frequency putative predictions (grey density), indicating that higher-frequency calls are strongly enriched for *bona fide* modification sites.

**Figure 2.**
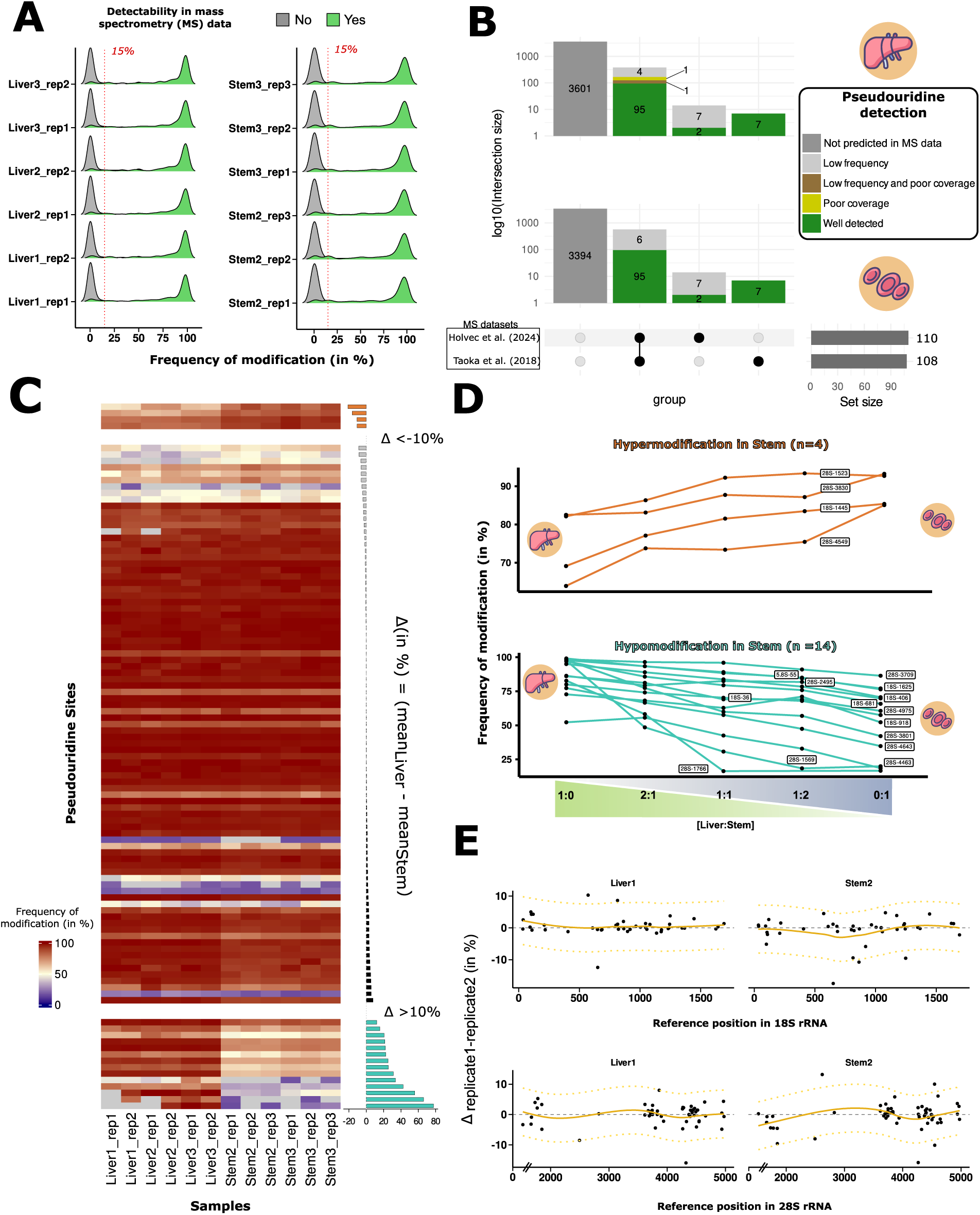
Nanopore sequencing precisely detects ribosomal RNA pseudouridine modifications. **(A)** Pseudouridine (Ψ) modification frequency distributions from nanopore sequencing in human liver (left) and hIPSC (right) samples. Mass-spectrometry-verified sites are green; unverified sites are grey. The red dotted line marks the 15% cutoff for high confidence pseudouridylated sites.**(B)** Classification of Ψ sites in liver (top) and hIPSC (bottom), showing detection confidence and overlap with published mass spectrometry datasets. The upset plot displays site counts categorized by presence/absence in two reference datasets (Holvec et al., 2024; Taoka et al., 2018), modification frequency, and coverage. Intersection colors indicate detection metrics per the legend. Thresholds: 15% for low frequency, 20 reads for lower coverage. **(C)** Context-dependent Ψ modification variation between liver and hIPSC. The heatmap shows nanopore-derived frequencies for MS-validated sites (rows) across replicates (columns). The right bar plot displays mean frequency difference (Δ = mean_liver − mean_stem) per site. Sites with |Δ| > 10%, indicating tissue/cell-type specificity, are emphasized. **(D)** Nanopore sequencing responsiveness to graded Ψ shifts in RNA mixtures. Liver and stem-cell RNA were mixed in varying proportions with increasing hIPSC fraction (x-axis). Modification frequencies for context-specific sites (panel C) were determined for each mixture. **(E)** Inter-replicate reproducibility of nanopore Ψ frequency estimation. Bland–Altman plots show technical replicate differences for each Ψ site in 18S rRNA (top) and 28S rRNA (bottom) in liver and stem cells. Points represent sites plotted by coordinate (x-axis) and frequency difference (Δ = replicate 1 − replicate 2; y-axis). Orange trend line and yellow confidence intervals show overall reproducibility.

To determine whether this separation reflected limited sequencing depth rather than biological signal, we analyzed previously generated direct RNA nanopore sequencing data from SKOV3ip1 human cell line rRNA (Supplementary Figure 1). Subsampling analyses revealed that increasing read depth caused many high-frequency putative predictions to shift toward the 15% threshold, consistent with coverage-dependent noise rather than true modification events. This behavior aligns with prior reports describing elevated false-positive rates in nanopore-based RNA modification detection and supports the current limitation of nanopore sequencing for de novo pseudouridine site discovery with existing basecalling models [42].

Based on these observations and on the literature, we defined conservative detection criteria requiring both a minimum local coverage of 20 reads [43] and an estimated modification frequency of at least 15%. Under these conditions, nanopore sequencing robustly detected 95 of the commonly MS-validated pseudouridine sites (Figure 2B). An additional seven sites reported by Taoka et al. [7] were also detected, whereas only two of the nine sites uniquely described by Holvec et al. [8] met these thresholds. Importantly, MS-validated sites with consistently low estimated frequencies (<15%) were not recovered even at increased coverage (Supplementary Figure 1B), indicating genuinely low stoichiometry that remains indistinguishable from background signal. Together, these results establish that nanopore-based pseudouridine detection in rRNA is reliable when constrained to validated sites and conservative frequency thresholds, providing a robust foundation for quantitative stoichiometry analyses presented below.

Having established conservative criteria for robust pseudouridine detection, we next assessed whether rRNA sequence variation influences nanopore-based stoichiometry estimates. Using the MS-confirmed pseudouridine sites, we indexed validated positions across five annotated structural variants of human 28S rRNA, enabling site-specific comparisons between variants. Because the 28SN5 variant (NCBI Gene ID: 100008589) is most commonly used as a reference in the literature, we compared pseudouridylation frequencies measured on 28SN5 to a weighted average of calls obtained from the remaining 28S variants. This analysis revealed no systematic differences in estimated modification levels across variants (Supplementary Figure 2), indicating that rRNA sequence heterogeneity does not substantially bias population-level stoichiometry estimates obtained by nanopore sequencing. Although this analysis does not exclude the possibility of variant-specific modification patterns at the single-molecule level, it supports the use of 28SN5 as a representative reference for subsequent analyses. In contrast to 28S rRNA, a single 18S variant (SN2) accounted for more than 90% of aligned reads across samples; this variant was therefore selected for all downstream 18S analyses. Together, these observations indicate that rRNA sequence variation does not confound nanopore-based pseudouridine quantification at the population level under the conditions examined.

### Nanopore sequencing maintains sensitivity across subtle changes in rRNA pseudouridylation stoichiometry

With a validated rRNA reference set and defined pseudouridine positions in place, we next examined whether direct RNA nanopore sequencing maintains quantitative sensitivity across subtle differences in pseudouridylation between biological contexts. We compared pseudouridine stoichiometries across all validated rRNA sites between human liver tissue and human induced pluripotent stem cells (hIPSCs) and ranked sites by their mean difference in modification levels (Figure 2C). This analysis identified a subset of 18 sites displaying pronounced context-dependent variation (absolute difference ≥ 0.1), including four sites that were relatively hyperpseudouridylated in stem cells and fourteen sites that were hyperpseudouridylated in liver samples.

To directly test sensitivity across diminishing differences in modification levels, we generated a series of mixtures combining liver and stem cell RNA at defined ratios, followed by library preparation, sequencing, and analysis using the same nanopore pipeline. For all 18 sites, estimated pseudouridylation levels changed progressively and monotonically with the mixing proportions (Figure 2D), demonstrating that nanopore-based measurements remain responsive even as differences between samples narrow. Both stem cell–enriched and liver-enriched sites exhibited gradual and reciprocal shifts as sample fractions were inverted.

As an illustrative example, pseudouridylation at site 18S-406 decreased from near-complete modification in liver-dominated mixtures to approximately 75% in stem cell–enriched samples, consistent with prior reports of rRNA epitranscriptomic remodeling during cellular differentiation [31,44]. Together, these results show that direct RNA nanopore sequencing maintains sensitivity across subtle changes in rRNA pseudouridylation stoichiometry, enabling quantitative tracking of rRNA modification dynamics across biological states.

### Nanopore sequencing provides reproducible and precise pseudouridylation estimates

To assess the reproducibility and precision of nanopore-based pseudouridylation measurements, we compared modification frequency estimates obtained from paired technical replicates. For each MS-validated pseudouridine site, replicate-to-replicate differences were calculated and visualized along the 18S and 28S rRNA sequences for representative liver and hIPSC samples (Figure 2E). Across both rRNA species and biological contexts, differences between replicates were tightly centered around zero, indicating strong agreement between independent sequencing runs. Although regions with dense pseudouridine content exhibited slightly increased variability, dispersion remained limited, with 95% confidence intervals ranging from ±4.8% to ±8.8% across samples. These results indicate that direct RNA nanopore sequencing provides stable and precise estimates of rRNA pseudouridylation stoichiometry when applied to a defined set of validated modification sites.

### Nanopore-based sequencing delivers robust, batch-independent quantification of rRNA pseudouridylation

To directly compare nanopore sequencing and HydraPsiSeq under matched experimental conditions, the same liver and hIPSC rRNA samples were analyzed using both approaches. Comparisons were restricted to pseudouridine positions supported by mass spectrometry (MS) and detected by both methods, yielding a common set of 107 (out of 117) rRNA sites used for downstream analyses (Figure 3A). Nanopore sequencing predictions made in either sample type were considered here. Absent from this common set were eight MS-validated sites from the 18S rRNA, excluded because they did not meet the nanopore detection thresholds defined above and because they were not represented in the predefined panel of rRNA loci selected a priori for PCR amplification in the HydraPsiSeq assay. Those sites are 18S-63, 18S-300, 18S-366, 18S-667, 18S-1003, 18S-1186, 18S-1360 and 18S-1596.

**Figure 3.**
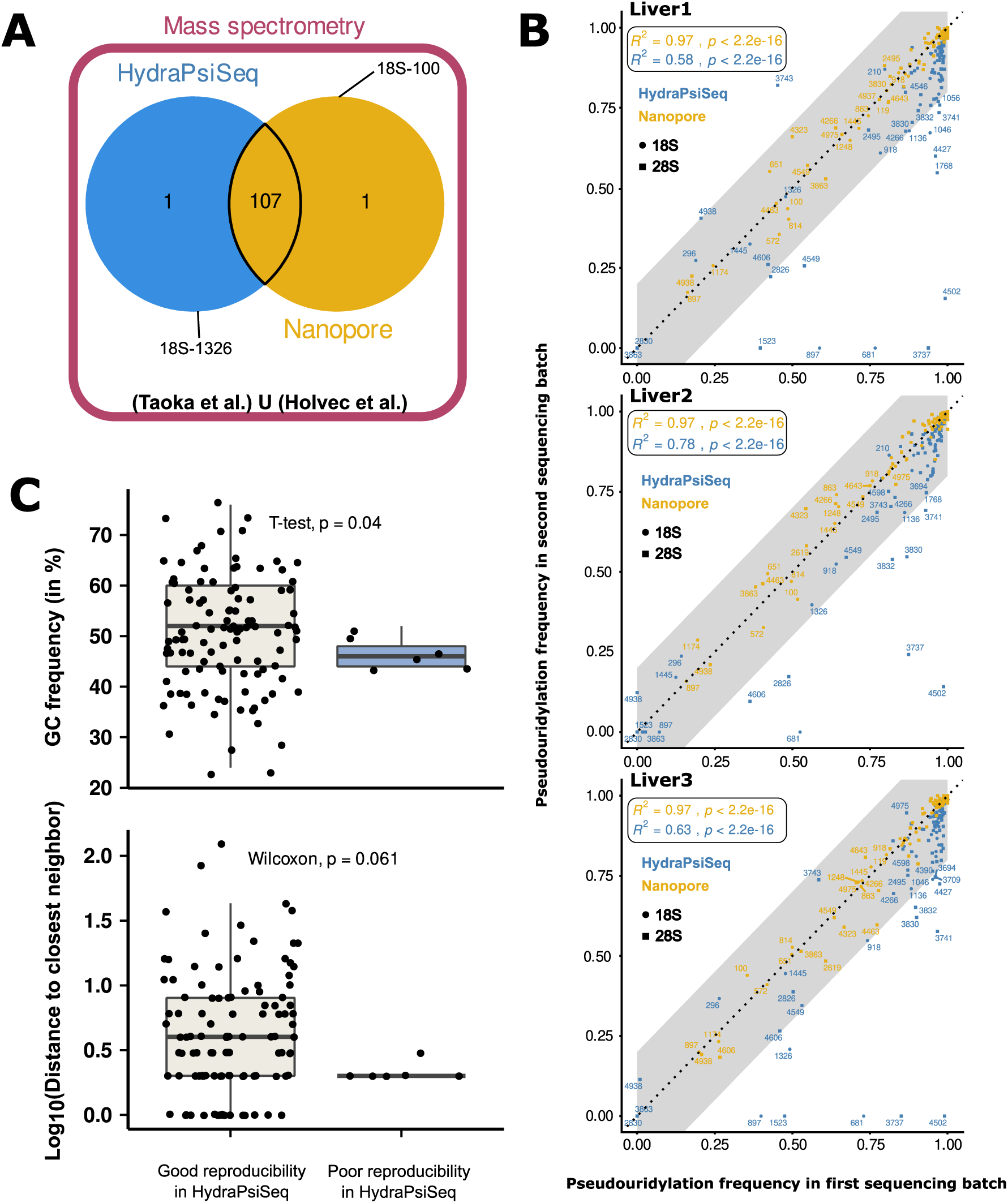
Reproducibility of pseudouridine (Ψ) detection across nanopore and HydraPsiSeq methods. (**A**) Intersection of pseudouridine sites detected by nanopore sequencing, HydraPsiSeq, and mass spectrometry. A Venn diagram illustrates the overlap among sites identified by the two sequencing-based methods and embedded in the union of both previously published mass spectrometry datasets. If in the union set but missing in either sequencing method, the sites are labeled by their rRNA coordinates for reference. **(B)** Batch-to-batch comparison of nanopore and HydraPsiSeq quantification. Scatter plots show pseudouridylation frequencies obtained from two independent sequencing batches for each of three liver samples (Liver1–3). For each method, the frequency estimates from batch 1 are plotted against that from batch 2. Nanopore-derived frequencies (gold/yellow) cluster tightly along the identity line, whereas HydraPsiSeq measurements (blue) show greater dispersion. The grey diagonal band represents a ±20% deviation threshold; points within this band exhibit high inter-batch concordance. The Pearson correlation is indicated in the upper left rectangle with an associated p-value for each pairwise comparison. **(C)** Sequence features associated with HydraPsiSeq reproducibility. Boxplot comparing surrounding GC-content (*Upper*) and log₁₀ distance to the nearest neighboring modified site (*Bottom*) for sites with good versus poor reproducibility in HydraPsiSeq (based on batch concordance in panel B). Individual sites are shown as points on both panels.

In addition, a small number of method-specific exceptions were observed. These included site 18S-1326, which carries an N1-methylpseudouridine and did not pass nanopore filtering, and site 18S-100, which was detected by nanopore sequencing but supported by only one of the two MS datasets. Together, these exclusions and exceptions reflect known challenges associated with sites exhibiting low stoichiometry or additional chemical modifications and define the set of rRNA positions amenable to robust cross-platform comparison.

To evaluate reproducibility across independent experimental batches, we next examined batch-level structure in the data. Principal component analysis revealed clear separation between samples of the different HydraPsiSeq batches alongside a component explaining 75% of the variance, indicating strong inter-batch effects (Supplementary Figure 3A). In contrast, nanopore-based estimates showed minimal batch-driven separation. This difference was further evident in site-by-site correlations across batches: nanopore sequencing produced highly consistent pseudouridylation estimates in both liver and hIPSC samples, whereas HydraPsiSeq displayed reduced reproducibility driven by null or highly variable calls at a subset of sites (Figure 3B; Supplementary Figures 3B and 3C).

Several HydraPsiSeq-affected sites, including 18S-681 and 28S-4502, deviated by more than 20% between batches and were flagged as poorly reproducible. Analysis of local sequence features revealed that these sites featured poor GC content and were frequently located within two nucleotides of another modified residue (Figure 3C). These characteristics are consistent with ambiguous cleavage patterns or reduced amplification efficiency and likely contribute to method-specific variability.

### Nanopore quantitative calling of pseudouridines is largely consistent with prior reported MS values

We next examined whether method-specific biases were associated with systematic differences in estimated modification levels. Pseudouridine fractions were averaged by method and compiled to evaluate site-by-site agreement in the measurements (Figure 4). Notably, while most predictions cluster near the identity line, precision varied substantially by method and site. For example, nanopore estimates with intermediate frequencies tended to show greater variability. This phenomenon is consistent with a known bias in the dorado model where local coverage drops for moderately modified sites, leading to lower precision in the quantification [45]. Extreme frequencies evaluated by HydraPsiSeq largely overlap with the poorly reproducible sites isolated in Figure 3B. Sites with discordant HydraPsiSeq estimates, located at the extremes of the plot, were consistently miscalled across samples and biological contexts, suggesting systematic capture inefficiencies driven by local sequence features rather than context-specific biological variation.

**Figure 4.**
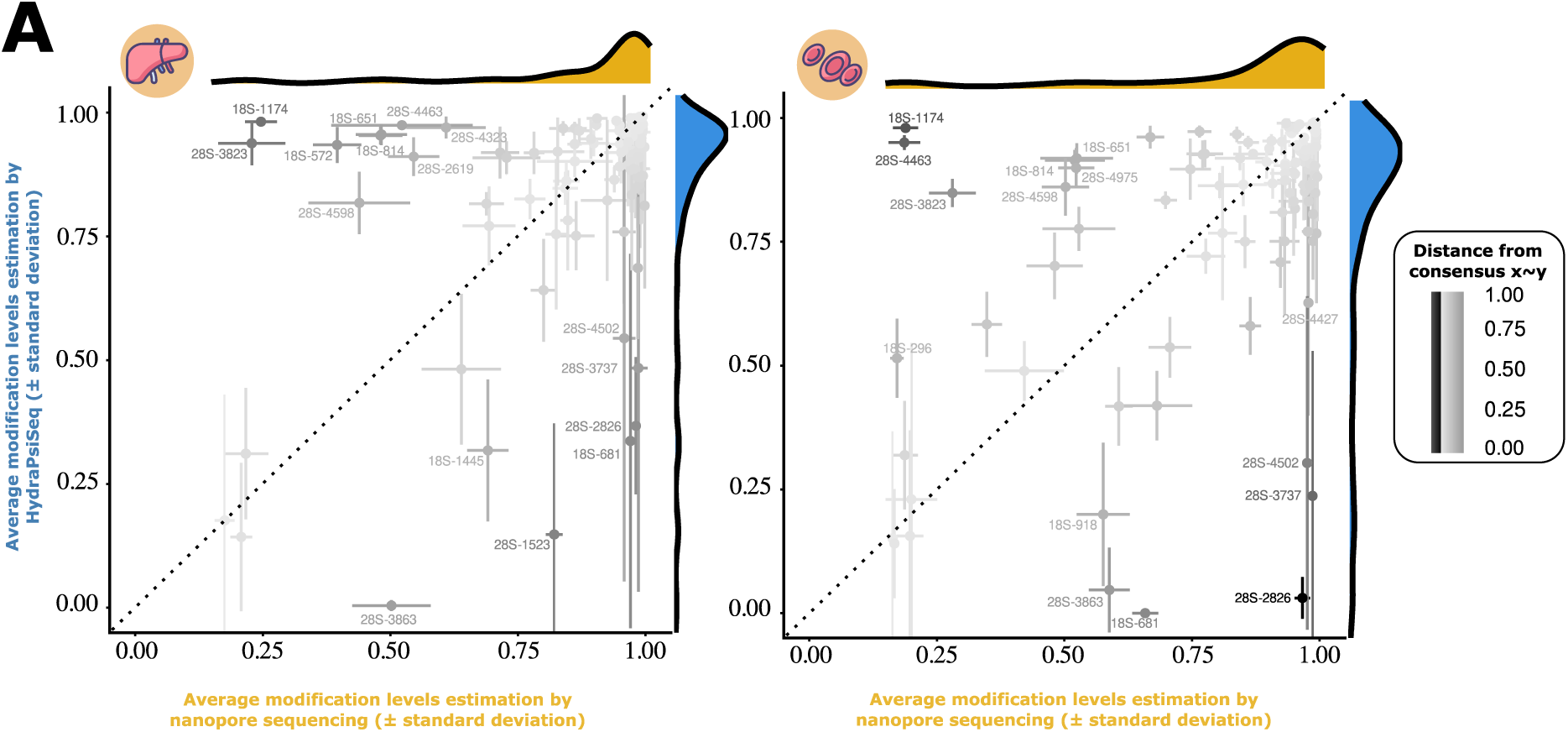
Pairwise cross-method concordance in liver and hIPSC samples. Comparison of per-site modification frequencies measured by nanopore sequencing (x-axis) and HydraPsiSeq (y-axis) across MS-validated Ψ sites. Points represent mean frequencies across replicates; horizontal and vertical error bars denote nanopore and HydraPsiSeq standard deviations. The diagonal line marks perfect concordance. Points are shaded by their distance from the concordance line, and marginal kernel density plots summarize the overall distributions for nanopore (top) and HydraPsiSeq (right). These plots highlight concordant sites as well as positions with substantial divergence between methods.

Overall, pseudouridine levels called by nanopore basecaller Dorado align with those of HydraPsiSeq for the most part. Although Dorado predictions exhibit relatively low variability, they conflict with HydraPsiSeq stoichiometry calls on about a third of rRNA sites with no way to adjudicate the discrepancies.

In an attempt to resolve the conflicts and to obtain a reliable quantitative estimation of pseudouridine levels for these regions, we extended the biological context to HeLa cells where previously published MS data have unambiguously characterized pseudouridine fractions. We then re-applied the current workflow to HeLa cells, performing nanopore direct RNA sequencing and HydraPsiSeq analysis as previously described [30]. For each sample, every site from the common set has its levels directly compared to the MS result (Figure 5A; see also Supplementary Figure 4A). Site-by-site Euclidian distance with MS frequencies confirmed existing monodirectional and bidirectional conflicts in quantification between MS and both high throughput approaches. If at least one methodology were disagreeing by 20% or more on the pseudouridine fraction, the site was further examined in Figure 5B. Estimates of the discordant sites by each of the three strategies demonstrated no unequivocal results, concentrating cases where MS agreed with one or the other approach. Closer to MS truth for fourteen sites, some nanopore direct RNA calls however fell off under our threshold of 15%. HydraPsiSeq PsiScore agreed more with MS for eleven rRNA sites, mostly due to large underestimations from Dorado. As for the remaining four observations, sequencing methods produced unconclusive values distant from MS estimations. This case-by-case partial agreement with distinct methods convinced us to explore the features that might hinder or nullify the quantitative capability of either approach. Driven by the relatively low GC richness of non-reproducible PsiScores, we mapped the rRNA locations that clash with the MS frequencies (Figure 5C). Surprisingly, pseudouridine sites in general were concentrated in low GC content regions. Discordant sites made no exception, seemingly located in AU rich valleys. Sites that HydraPsiSeq misquantifies on the 18S display a bimodal GC content pattern, with the poorly reproducible sites from Figure 3B predominantly found in the GC-poor peaks. Miscalled sites by nanopore even show underrepresented GC richness (Figure 5D). Additionally, the quality of dorado predictions appears to be lower for those same sites (Figure 5E). Lastly, we compiled the differences in prediction levels from the two methods in hIPSC, HeLa, and liver samples to isolate context-specific miscalls (Supplementary Figure 4B). It resulted in 73 rRNA sites being concordantly quantified and 5 sites constantly being conflictual across the three biological contexts possibly due to a specific local structure around these regions.

**Figure 5.**
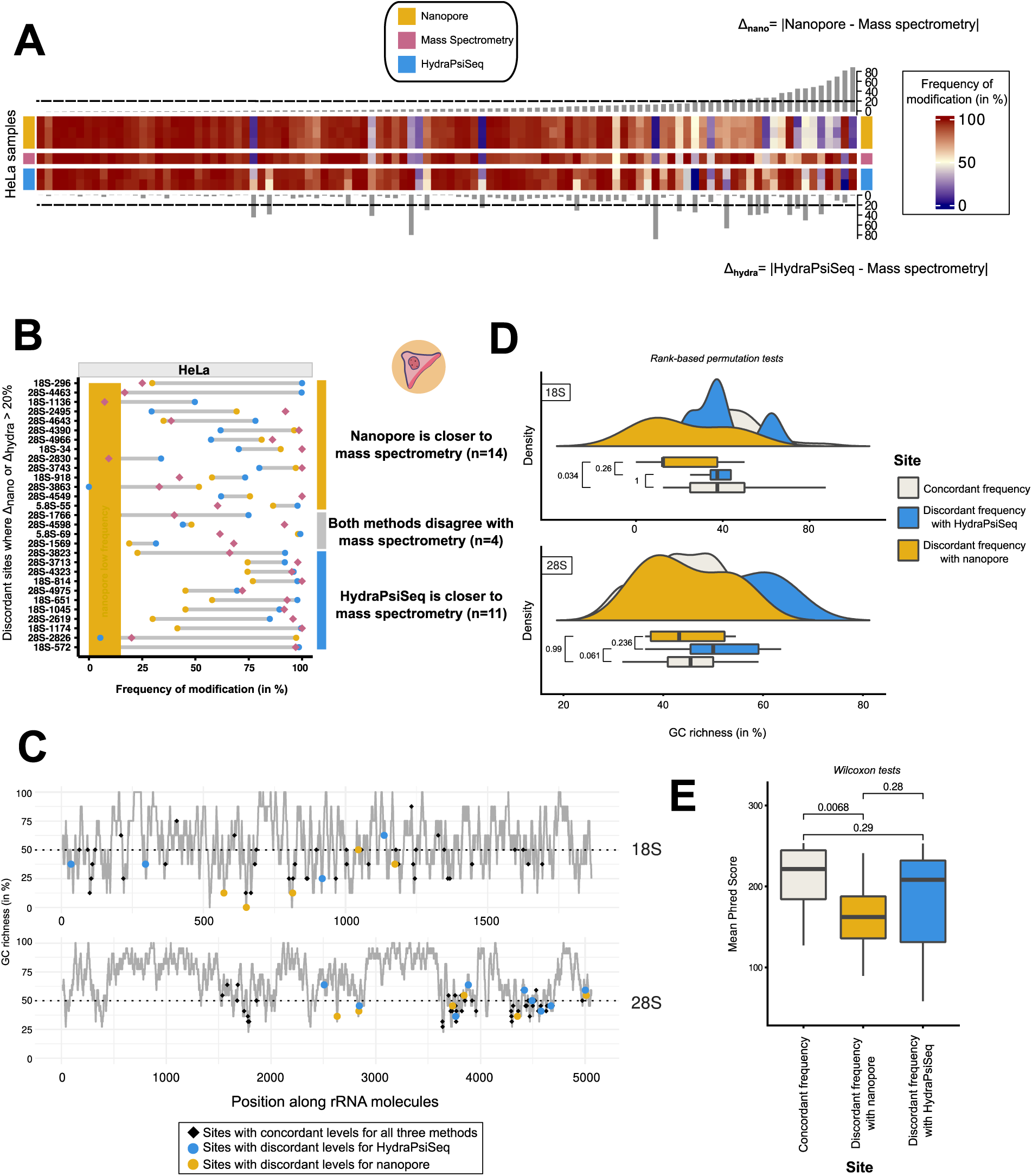
Resolution of method disagreements in HeLa cells and limitations partially explained by structural features. **(A)** Heatmap of the frequency of modification for rRNA sites (in columns) by MS, HydraPsiSeq or nanopore sequencing (in rows) done in HeLa cells. Top and bottom barplots indicate the distance between the fraction measured by MS and one high throughput method (top for nanopore; bottom for HydraPsiSeq) for each site. Columns are ordered by the nanopore-MS distance, and dashed lines mark when the distance goes over 20%. **(B)** Modification frequencies for all discordant sites across methods and sample types. Rows correspond to individual Ψ sites; frequencies from MS (pink diamonds), HydraPsiSeq (blue circles), and nanopore (gold circles) are shown for HeLa samples. A faded yellow tile demarcates the area where nanopore predictions fall under 15% of frequency. To the right, the color bar counts sites according to whether nanopore or HydraPsiSeq is closer to the MS reference. **(C)** Line plot accounts for the GC richness of 18S (top) and 28S (bottom) rRNA sites. Values are smoothed by a sliding window of 8 and 22 nucleotides for 18S and 28S molecules respectively. On the grey lines, pseudouridylated residues appear as small black diamonds while sites with miscalled fractions are decorated with circles colored after the method at fault (gold yellow for nanopore; blue for HydraPsiSeq). **(D)** Raincloud plots of 18S and 28S rRNA showing GC richness as kernel densities and boxplots for three subsets of sites: HydraPsiSeq and nanopore agree (white) or disagree (gold yellow for nanopore; blue for HydraPsiSeq). Each subset of sites is compared to the other with a ranked permutation test from which p-values on the left are originating. **(E)** Boxplots of the compiled and averaged Phred scores visualized for each subset and compared with pairwise Wilcoxon tests. Boxes are colored after the subset they represent.

Thus, leveraging MS stoichiometries in HeLa cells made the resolution of most conflicting sites possible and confirmed the ability of basecaller dorado to rightly quantify pseudouridine fractions on 92 rRNA sites in HeLa. This benchmark revealed the proficiency of the current nanopore basecalling model for accurately characterizing 85% pseudouridine stoichiometries of MS expected human rRNA sites across vastly different contexts, leaving only 15 sites with unreliable predictions.

## Discussion

Lately, post-transcriptional RNA modifications are increasingly recognized for their dynamic transcriptome-wide changes. Sub-stoichiometric levels on key RNA structures can mark nuanced control over gene expression, leveraging RNA stability. In that direction, tRNA hypomodifications distinguish marginal zone B cells from neuroblastoma cells [46], while pseudouridine in pre-mRNA directly modulates splicing [47]. This new layer remains relatively uncharted despite its potential for gene expression regulation. This is also true for the densely modified human rRNA where modifications equilibrium impact remains unresolved. Yet evidence about the role of the epitranscriptome in the modeling of the ribosome from its biogenesis to its translation program is accumulating [8,11,48]. As resolution of methodologies is improving, post-transcriptional modifications emerge as a tangible layer of ribosome heterogeneity. In that view, several high throughput methods have tried to tackle the challenge of pseudouridine variability in the densely modified ribosomal RNA.

Among them, nanopore sequencing exploits the distinct current disruption signals produced by RNA modifications, enabling their detection using modification-aware basecalling models. Nonetheless, validations of these predictions in non-artificial conditions are severely lacking, especially in densely modified RNAs such as in ribosomal RNA. By examining rRNA sites case-by-case in liver tissues and hIPSCs, this study investigates the capabilities of nanopore based sequencing method for pseudouridine quantification over the chemically reactive short read sequencing approach HydraPsiSeq.

We started with the evaluation of nanopore’s ability to capture a substantive landscape of pseudouridines, relying on an extended repertoire of mass spectrometry-validated positions along the RNA. Although MS might be the closest to a gold standard method in pseudouridine detection, it is likely that there are pseudouridine positions still missing outside the union of the MS datasets and therefore not considered later in the study. In that sense, nanopore calls a significant number of sites (∼1400) as highly modified in Figure 2A that are listed as not detected in MS (grey bulge near 100% frequency). This number drops substantially when considering that predictions must be seen in at least half of the samples and further than 3 nucleotides from already known pseudouridine or methylated uridine with a total of 34 candidates. With deeper coverage in Supplementary Figure 1A this population shrinks but does not disappear, suggesting that nanopore sequencing can detect those putative modifications. Further studies will be required to determine whether these sites are truly modified.

Since rRNA pseudouridines may serve as tissue-specific epitranscriptomic signatures [31], sites with typically low modification levels may show substantially increased stoichiometry in specific biological contexts. This is the case for site 1766 of the 28S rRNA that falls under a critical threshold of 17% in stem cells on Figure 2D while being rather highly pseudouridylated at 82% in liver samples. Sequencing a greater diversity of tissues and conditions will enable to document the range of modification levels of each site and might provide further evidence of the physiological relevance of currently unannotated sites.

Sequence heterogeneity among rRNA copies may also contribute to underestimating pseudouridine site’s stoichiometry. Here, we minimize the impact of sequence variability by rejecting reads that do not align with prevalent copies of 28S and 18S rRNA. Even though rRNA sequence variation does not alter overall modification stoichiometry as shown in Supplementary Figure 2, it could affect local RNA structure and accessibility, thereby influencing which sites can be modified [14].

While rRNA copy variants consist primarily of indels, no studies have systematically examined pseudouridine patterns at single-molecule resolution. This new layer of analysis could unveil whether specific rRNA sequence variants are associated with distinct modification patterns, potentially linking sequence variations to co-regulated post-transcriptional modification events. Because modification callers provide base-level confidence scores, nanopore sequencing now enables increasingly feasible single-molecule analysis. These analyses could shed light on core mechanisms of stoichiometric balance.

Our study featured the comparison of nanopore sequencing and HydraPsiSeq contrasting holistic and deterministic approaches for pseudouridines quantification. A key distinction is that the current basecalling model generates unsupervised predictions on modified nucleotides without prior knowledge of modification sites, whereas HydraPsiSeq targets a predefined list of known rRNA locations. This difference allows nanopore sequencing to detect a broader number of putative sites. The exclusive sites labelled in Figure 3A are quite illustrative of this phenomenon: 18S-1326 is missed by nanopore sequencing probably because it carries an additional methylation however one can assume that the upcoming models will eventually capture this site with good metrics. On the other hand, 18S-100 is not included as a target in HydraPsiSeq and as such cannot be quantified. Furthermore, in nanopore sequencing, the acquisition of electric signals for each nucleotide and the calling of post-transcriptional modifications are two distinct steps. Therefore, the per-base current intensity, stored in .pod5 files, can be exploited *a posteriori* with the latest models. The virtually endless lifetime of nanopore data overcomes the one-time capture of pseudouridines by HydraPsiSeq. All the more so, as the novel generation of models now call multiple modifications simultaneously and are expected to eventually broaden their scope to many canonical post-transcriptional modifications reported on the RNA. As mentioned in the methods section, for this manuscript, we tried basecalling using models 5.1, 5.2 and the most recent version 5.3, the two latter supporting detection of additional RNA modifications. The tradeoff was a loss of precision for pseudouridine (see Supplementary Figure 5A-E, shown for 5.1 vs 5.3). After testing the model against the MS values in HeLa cells, we observed a general leveling down of pseudouridine for model 5.3 compared to model 5.1. By being agnostic to neighbouring modifications, model 5.1 seems to gain stability but might still miscall pseudouridines despite the stringent precautions adopted in this paper. So for now, the model limiting the diversity of modifications detected performs best, but it is expected that with time predictions for simultaneous detection of multiple types of modifications will continue to improve, and the analyses of the nanopore datasets presented here can be repeated with the new models, ever increasing their accuracy.

In addition to the lack of re-usability of its datasets when future improvements of the technology are introduced, HydraPsiSeq also suffers from vulnerability to batch variations, displaying contradictory PsiScores sites in two runs. The systematic error of a few sites in Figure 3B has led us to flag their lower GC content and a higher number of proximal modified neighbors in Figure 3C. Lot-to-lot variation in reactivity and hydrazine ages may have contributed to these discrepancies, as different reagent batches were used across experimental runs. Nevertheless, this highlights a limitation of the method: it can introduce substantial technical bias that is difficult to control. Moreover, it seems that hydrazine has a partial reactivity with methylated uridine [49].

Comparison of the quantification between nanopore sequencing and HydraPsiSeq revealed discordance on roughly 15% of the MS validated sites. While some discrepancies can be attributed to poor reproducibility of HydraPsiSeq, other errors look less categorical with intermediate frequency predicted. For these sites, local structures may, each in their specific way, impair one methodology or the other. Prior work using the Guppy basecaller suggested that context sequence deeply influences the decision to call a modification lengthening the k-mer window up to +/− 5 nucleotides [50]. For instance, the presence of guanines upstream of the queried sequence is associated with slower translocation through the pore, resulting in an altered ionic current signature. In Figure 5C, 5D, and 5E, we point out a significant link between the low GC content and weaker quality in regions where nanopore basecaller miscalls pseudouridines.

Moving to HeLa cells made it possible to resolve the frequency of all but four discordant sites by bringing MS data in. Of the sites miscalled and apart from site 28S-2826, nanopore model 5.1 constantly underestimated pseudouridine abundance. This behavior was consistently observed during testing with the expanded model 5.3, suggesting that densely structured environments may interfere with ionic current signaling within the pore, resulting in downward levels of prediction.

We finally summarize and count the wrongly estimated objects in Supplementary Figure 4B and put the emphasis on a contentious subset constant across biological contexts. These five sites, all of which emerged evolutionarily in high eukaryotes, may necessitate future investigations to identify systematic features that explain why they escape clear-cut quantification by sequencing approaches.

Beyond the specific biological comparisons presented, the workflow described here offers practical advantages for researchers seeking to profile rRNA pseudouridylation across large sample cohorts or clinically derived material. The low cost per sample (∼$60 with 96 barcodes), short library preparation time, and reusability of raw signal data with future basecalling models collectively lower the barrier for epitranscriptomic profiling in settings where mass spectrometry infrastructure is inaccessible or where sample availability limits input material. The defined detection thresholds established here, a minimum modification frequency of 15% and local coverage of at least 20 reads, provide a ready-to-use analytical framework that can be directly adopted by groups profiling pseudouridylation in patient-derived tissues, primary cells, or rare biological samples. As rRNA modification stoichiometry continues to emerge as a biologically and clinically relevant phenotype, the scalability and reproducibility demonstrated in this study position nanopore direct RNA sequencing as a practical tool for population-scale epitranscriptomic studies.

## Conclusion

This nanopore-based pipeline demonstrated performance comparable to established orthogonal methods such as mass spectrometry for quantitative assessment of pseudouridine fractions in human rRNA. The method recovered a major part of previously annotated rRNA pseudouridines and estimated their relative abundance consistently across independent batches of sequencing, biological contexts, and mixed samples. Taken together, the scalability of the approach, the short preparation and processing time, the high exploitability of the resulting data, and the possibility of performing multiple modification calls indicate that nanopore sequencing is a strong candidate to become a routine method for epitranscriptomic studies (see Table 1). Although it marks a milestone toward a comprehensive appreciation of the human post-transcriptional landscape, this nanopore approach remains dependent on orthogonal strategies for a subset of rRNA regions, notably featuring low GC richness and for *de novo* predictions. It is yet undeniable that nanopore-derived strategies will be of great help in the future. The advances with long-read sequencing continue to expand our analytical resolution, while methods integrating epitranscriptomic profiles and other features offer considerable promises to tackle mechanistic challenges and ultimately translation-associated diseases.

**Table 1.** Summary of the pros and cons of methods used in this benchmarking study.

|  | **Mass Spectrometry** | **HydraPsiSeq** | **Nanopore** |
| --- | --- | --- | --- |
| **Scalability** | Poor (low throughput) | Good | Good (up to 96 barcodes for demultiplexing) |
| **Specificity to pseudouridine** | Good | Poor (indirect, through hydrazine reactivity) | Good |
| **Sample Reproducibility** | Good (with standardized conditions) | Moderate (strong batch effects) | Good |
| **Suitable for de novo site exploration** | Yes | No | Potentially (with adequate training data) |
| **Cost per sample (consumables and library preparation included)** | ~200\$ | ~150\$ | ~60\$ with 96 barcodes |
| **Quantification accuracy** | Very good (absolute stoichiometry with standards) | Moderate (relative quantification, requires normalization) | Good (per-read modification probability) |
| **Detection biases** | Completeness of RNA digestion | Determined list of targeted sites<br>Influence of nearby modified sites | Unknown model training bias<br>Influence of local context |
| **RNA input required** | 1µg of RNA | 200 ng of RNA | 500 ng of polyadenylated RNA |
| **Turnover** | Slow (1-3 days for LC-MS/MS run + analysis) | Moderate (~1-2 days sequencing+analysis) | Fast (<1 day sequencing+analysis) |
| **Reusability** | One time | One time | Re-Analyzable with future models |

## Resource availability

### Lead contact

Requests for further information and resources should be directed to and will be fulfilled by the lead contact, Michelle Scott.

### Material availability

This study did not generate new unique reagents.

### Date and code availability

Converted fast5 files obtained from the nanopore direct RNA sequencing runs and processed HydraPsiSeq PsiScore tables have been deposited in the European Nucleotide Archive (ENA) at EMBL-EBI under accession number PRJEB108556. The pipeline used for the processing of the nanopore data is available at https://github.com/Baud-de-Preval/rRNA-PsiPore.

## Methods

### Acquisition of rRNA liver samples

Liver RNA samples were purchased from BioChain, as previously described [51] (RNA integrity was assessed using the 2100 Bioanalyzer (Agilent Technologies, USA).

### Culture of SKOV3ip1 and HeLa cell lines

The SKOV3ip1 human ovarian carcinoma cells were cultured in DMEM/F12 (50:50) medium supplemented with 10% fetal bovine serum (FBS) and 2 mM L-glutamine. HeLa human cervical cancer cells were cultured in DMEM medium supplemented with 10% FBS with 1% MEM non-essential amino acids. Cells were maintained at 37°C in a humidified incubator with 5% CO₂ and were periodically tested for mycoplasma contamination.

### Isolation of human induced pluripotent stem cells (hIPSC)

Human induced pluripotent stem cells (iPSCs) were maintained under feeder-free conditions on Cell Basement Membrane-coated dishes (150 μg/mL in DMEM:F-12, 2 mL per 6 cm² dish, incubated 1 hour at 37°C). Cryopreserved cells were thawed rapidly (1–2 minutes in 37°C water bath), transferred dropwise into Pluripotent Stem Cell SFM XF/FF medium, centrifuged at 200 × g for 5 minutes, and resuspended in medium supplemented with 10 μM ROCK Inhibitor Y27632. Cell aggregates were seeded onto prepared dishes and incubated at 37°C with 5% CO₂. A complete medium change was performed daily, with non-adherent cells removed on day 1 post-thaw. Cultures were monitored microscopically for compact colonies with high nucleus-to-cytoplasm ratios and prominent nucleoli; differentiated cells (<10% of culture) were aspirated using a fine-tipped pipette. Cultures were passaged at 80% confluence (every 4–5 days) using Stem Cell Dissociation Reagent (0.5 U/mL, 10–15 minutes at 37°C) until colony edges loosened, followed by gentle rinsing and trituration in medium containing ROCK inhibitor to maintain cell aggregates (not single cells). Cells were centrifuged, resuspended, and replated at a 1:4 split ratio onto fresh matrix-coated dishes; subsequent daily medium changes omitted ROCK inhibitor.

### RNA Extraction Protocol

Total RNA was extracted from SKOV3ip1 and HeLa cells as previously described [15]. Total RNA was extracted from stem cells using the RNeasy Mini Kit according to the manufacturer’s quick-start protocol. Cell pellets (≤1 × 10⁷ cells) were lysed in Buffer RLT (600 μL for 1 × 10⁷ cells) with addition of 10 μL β-mercaptoethanol per 1 mL buffer to inhibit RNase activity, followed by homogenization using vortex. Lysates were centrifuged at maximum speed for 3 minutes, and the supernatant was transferred to fresh tubes. One volume of 70% ethanol was added to the lysate and mixed thoroughly without centrifugation. Up to 700 μL of the lysate-ethanol mixture was applied to RNeasy Mini spin columns in 2 mL collection tubes and centrifuged at 13000 × g for 15 seconds, with flow-through discarded. Sequential wash steps were performed by adding 700 μL Buffer RW1 (15 s, 13000 × g), followed by two washes with 500 μL Buffer RPE (first wash: 15 s; second wash: 2 min at ≥8000 × g). An optional drying step was performed by centrifuging the column at full speed for 1 minute to dry the membrane. RNA was eluted by adding 50 μL RNase-free water directly to the spin column membrane, centrifuging for 1 minute at 13000 × g.

### Polyadenylation of RNAs

PolyA tailing reactions were performed as previously described [52] for liver, stem cell, HeLa and SKOV3ip1 RNA samples. Briefly, 500ng of total RNA was diluted in 10µL of nuclease-free water and heated at 80°C for 2 minutes. 10µL of the reaction mixture was added to each sample, which contained 2µL of NEB poly(A) buffer (NEB, B0276S), 2µL of an in-house 10X poly(A) buffer (containing 1μl of 1 M MnCl2 (Sigma, M1787), 4μl of 100 mM DTT, 4μl of 50 mg/ml BSA (Life Technologies, AM2616), and 31μl of nuclease-free water for final concentrations of 25 mM MnCl2, 10 mM DTT, and 5 mg/ml BSA), 1µL 10 mM ATP, 1µL SUPERase.in (20 U/µL, Life Technologies, AM2694), 1µL Clontech poly(A) polymerase (Takara Bio, 2180B) and 3µL of nuclease-free water. This resulted in final concentrations of 1X NEB poly(A) buffer, 1x in house buffer, 0.5 mM ATP, and 1U/µL of SUPERase.in. Reactions were incubated at 37°C for 7.5 minutes and immediately stopped by adding 0.5µL of 500 mM EDTA and placed on ice. RNA was purified using Zymo RNA Clean and Concentrator columns (Zymo Research, R1016).

### Library preparation for Nanopore RNA direct sequencing

The following preparation was performed at the RNomics Platform of the Université de Sherbrooke. Preparation of the libraries was performed according to the manufacturer’s instructions (direct_RNA_sequencing_SQK-RNA004_DRS_9195_v4_revJ_24Nov2025-promethion), with following exception: 24 barcodes [39] were ligated to 500ng of RNA instead of the RTA, and samples were subsequently pooled after the RT reaction. Libraries were then loaded on a FLO-PRO004RA flowcell and sequenced on a PromethION 2 Solo using MinKNOW software.

### Base-calling and alignment of direct RNA sequencing

pod5 files were basecalled with native Oxford Nanopore Technologies basecaller *dorado* v*1.1.1* [35] jointly with the modification-aware models rna004_130bps_sup v5.3, rna004_130bps_hac v5.3, rna004_130bps_supp v5.2 for *pseU_2meU, and* rna004_130bps_sup for *pseU* v5.1 to compare their performance. The latter showed stronger reproducibility on sequencing replicates and a lower number of predicted novel pseudouridine sites outside the list of validated MS sites (see Supplementary Figure 5). Collectively, these elements supported the selection of model 5.1 in preference to model 5.3. Reads were aligned using *Minimap2* v2.28 [53] with the parameters “-x map-ont -N 0 -k 13” against five rRNA sequences for the 28S, 2 rRNA sequences for the 18S and one sequence for the 5.8S and 5S. Variants were selected from the representative copies defined in RefSeq : 28SN1 to 28SN5 for the 28S (respective NCBI Gene IDs: 106632264, 109864282, 109910382, 109864272 and 100008589), 18SN1 to 18SN5 for the 18S (respective NCBI Gene IDs: 106631781, 109864280, 109910380, 109864273 and 106631781). Perfect sequence identity led us to merge the different 18S variants into 18SN1-5 and 18SN2-3-4.

### Demultiplexing direct RNA sequencing

*Seqtagger* v1.0d [39] was used to reattribute reads to their corresponding samples through 24 DNA barcodes introduced during library preparation. Index file *RNA004.demux.tsv.gz* was built and bam files were demultiplexed with *bam_split_by_barcode.py*. For each sequencing run, this step resulted in the creation of 96 bam sub-files from which only the first 24 were kept.

### Compilation of per-base predictions

Indexation and sorting of the bam files was done using *samtools* v1.22.1 [54]. The files were then fed to *modkit* v0.5.1 [36] to extract and to compile pseudouridylation events across each uridine base with the options “--motif T 0 --filter-threshold T:0.95”.

### Analysis of basecalled pseudouridines

All of the statistical analyses were done on *R* v 4.5.2 using *ggplot* [55]*, ggpubr* [56]*, perm* [57]*, ggridges* [58]*, tidyverse* [59]*, ComplexHeatmap* [60]*, ComplexUpset* [61]*, pheatmap* [62]*, corrplot* [63]*, FactoMineR* [64] and *factoExtra* [65]. HydraPsiSeq and nanopore predictions were compared to the union of all pseudouridines identified by mass spectrometry in Holvec [8] or Taoka [7]. Nanopore predictions under 15% of frequency or less than 20 reads passing quality control for local coverage were discarded. HydraPsiSeq’s poorly reproducible sites were defined as such if seen at least twice outside the 20% deviation area around the identity line in the liver sample. Pseudouridine ratios for MS data in HeLa cells were taken from Taoka et al [7]. We used descriptive statistics of estimates in HeLa between mass spectrometry and sequencing methods to introduce the following condition for discordant sites:

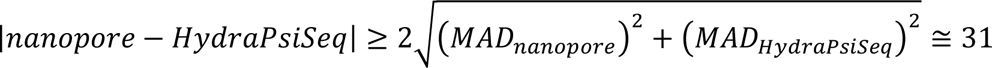 with MAD being short for mean absolute difference.

### Feature extraction of the rRNA sites

A list of modified nucleotides in rRNA was produced using the MS datasets. Using the *Clustal Omega* [66] multiple sequence aligner, shifts in the base numbering were inferred for each modified nucleotide across each variant (see Supplementary Table 1). Distance to the closest modified neighbor was computed as the shortest distance needed to meet another modified base. GC richness was calculated using the python module *BioSeqUtils* [67] on a sliding window proportional to the length of each molecule: 22 nucleotides for variant 28SN5 and 8 nucleotides for 18SN2-4. Phred scores were obtained from the bam files using samtools v1.22.1 with the command: “*samtools mpileup ${input} -f reference.fa -M”* and then averaged for each uridine along rRNA molecules. Evolutionary conservation of the sites was pulled from Taoka et al. on Supplementary Table S9-1 and S9-2.

### HydraPsiSeq sample preparation and sequencing

Total RNA (50-300 ng) from different human cells or tissues was subjected to hydrazine treatment (50% final concentration) for 45 min on ice. The reaction was stopped by ethanol precipitation using 0.3M NaOAc, pH5.2 and Glycoblue and incubated 30 min at −80°C. After centrifugation, the pellet was washed twice with 80% ethanol and resuspended in 1M aniline, pH4.5. The reaction was incubated in dark for 15 min at 60°C and processed as described above, by ethanol precipitation.

RNA fragments obtained by hydrazine/aniline treatment were dephosphorylated at the 3’-end for 6h at 37°C using 10 U of T4 PNK in 100 mM Tris-HCl pH6.5, 100 mM MgOAc and 5 mM β-mercaptoethanol. T4 PNK was inactivated by incubation for 20 min at 65°C. RNA was extracted by phenol:chloroform and ethanol precipitated. The pellet was resuspended in RNase free water. 3’-dephosphorylated RNA fragments were converted to library using the NEBNext® Small RNA Library Prep Set for Illumina® (NEB ref E7330S, USA) following the manufacturer’s recommendations. DNA library was quantified using a fluorometer (Qubit 3.0 fluorometer, Invitrogen, USA) and qualified using a High Sensitivity DNA chip on Agilent Bioanalyzer 2100. Libraries were multiplexed and subjected for high-throughput sequencing on an Illumina NextSeq2000 instrument with a 50 bp single-end read mode.

### HydraPsiSeq computational pipeline

High quality raw sequencing reads (> Q30) were subjected to trimming using Trimmomatic v0.39 [68] with the following parameters: MINLEN:08, STRINGENCY:7, AVGQUAL:30, trimmed reads were further aligned to the human rRNA (or rRNA/tRNA) reference sequence (acc. n° if available) using bowtie2 v2.4.4 [69] in end-to-end mode (--no-unal --no-1mm-upfront -D 15 -R 2 -N 0 -L 10 -i S,1,1.15 as other bowtie2 parameters), only uniquely mapped reads in positive orientation were retained for further analysis. Mapped and sorted *.bam file was transformed into *.bed format. Locations of 5’-extremities of mapped unique reads were counted from *.bed file, giving raw cleavage profile. Reads’ 5’-end counts were normalized to local background in rolling window of 10 nucleotides; values for U residues were excluded to calculate the median. Locally normalized U profiles (NormUcount) were further transformed to U cleavage profiles only, by omitting values for other nucleotides. Resulting U profiles were used for calculation of “RiboMethSeq-like” scores (ScoreMEAN, A, B and PsiScore, equivalent to ScoreC in RiboMethSeq). Window of 4 neighboring nucleotides (+/-2 nt) was used for score calculation [70].

ScoreMEAN for each position is calculated in two steps, as follows: first, a ratio for number of cumulated 5’/3’-reads ends between preceding and following position is defined and, second, ScoreMEAN is calculated as a ratio of a drop for a given position compared to the average and variation for 4 neighboring positions (−2/+2). PsiScore(ScoreC2 in RiboMethSeq), is calculated using the following formula : PsiScore = 1-ni/(0.5*(SUM(nj*Wj)/SUM(Wj)+SUM(nk*Wk)/SUM(Wk)), where ni – cumulated 5’-end U count for a given position, j – varies from i-2 to i-1, k varies from i+1 to i+2, Weight parameters are defined as 1.0 for −1 /+1 and 0.9 for −2/+2 positions. Quantification of Psi residues was done using PsiScore (similar to scoreC2 used in RiboMethSeq [70,71]). This score is used for quantification of the modification level since it keeps linear calibration curve between protection and molar ratio of Psi residue in RNA.

## Supplementary Information

**Additional file 1**: Supplementary figures

## Abbreviations

cDNA: complementary DNA
CMC: N-cyclohexyl-N′-β-(4-methylmorpholinium)ethylcarbodiimide
DRS: Direct RNA sequencing
hIPSC: human Induced Pluripotent Stem Cell
MS: mass spectrometry
rRNA: ribosomal RNA

## Declarations

### Ethics approval and consent to participate

Not applicable

### Consent for publication

Not applicable

### Declaration of interests

The authors declare that they have no competing interests.

### Funding

This work was supported by Canadian institute of health research (CIHR) grants to M.S.S. and S.A.E. (PJT 479838 and 486207) and Cancer Research Society (CRS) grant (25108) by NT. LFG is supported by a Fonds de Recherche du Québec – Santé (FRQS) doctoral scholarship. M.S.S. holds the Canada Research Chair in Bioinformatics of Non-Coding RNA and S.A.E. holds the Canada Research Chair in RNA Biology and Cancer Genomics.

PLN is supported by the Alberta Innovates Graduate Fellowship.

### Author contributions

BSP, MSS, and SAE conceived the study. BSP, MSS, LFG, and SAE designed the experiments, and BSP integrated the data including basecalling, read mapping and processing and other analysis. BSP made all the figures. LFG performed the RNA extraction and polyadenylation for all the nanopore samples; MD carried out nanopore library preparation and sequencing. VM and YM generated the HydraPsiSeq data for all samples. PLN and NT provided hIPSC RNA. BSP wrote the manuscript and iteratively improved it with MSS and SAE. All authors read and approved the final manuscript.

## Acknowledgements

The authors would like to thank Lauren Kwiatek and Anne-Marie Landry-Voyer from François Bachand’s laboratory at the Université de Sherbrooke for generously providing the HeLa cell line used in this study, the Digital Research Alliance of Canada for providing computing infrastructures and the Scott and Abou Elela groups for the helpful discussions.

## Notes

### Competing Interest Statement

The authors have declared no competing interest.

