## Supplementary figures for "Nanopore Direct RNA Sequencing Enables Reproducible, Site-Resolved Pseudouridine Quantification in Human Ribosomal RNA"

Baudouin S de Préval et al. 2026

-

*Supplementary Figures S1 to S5*

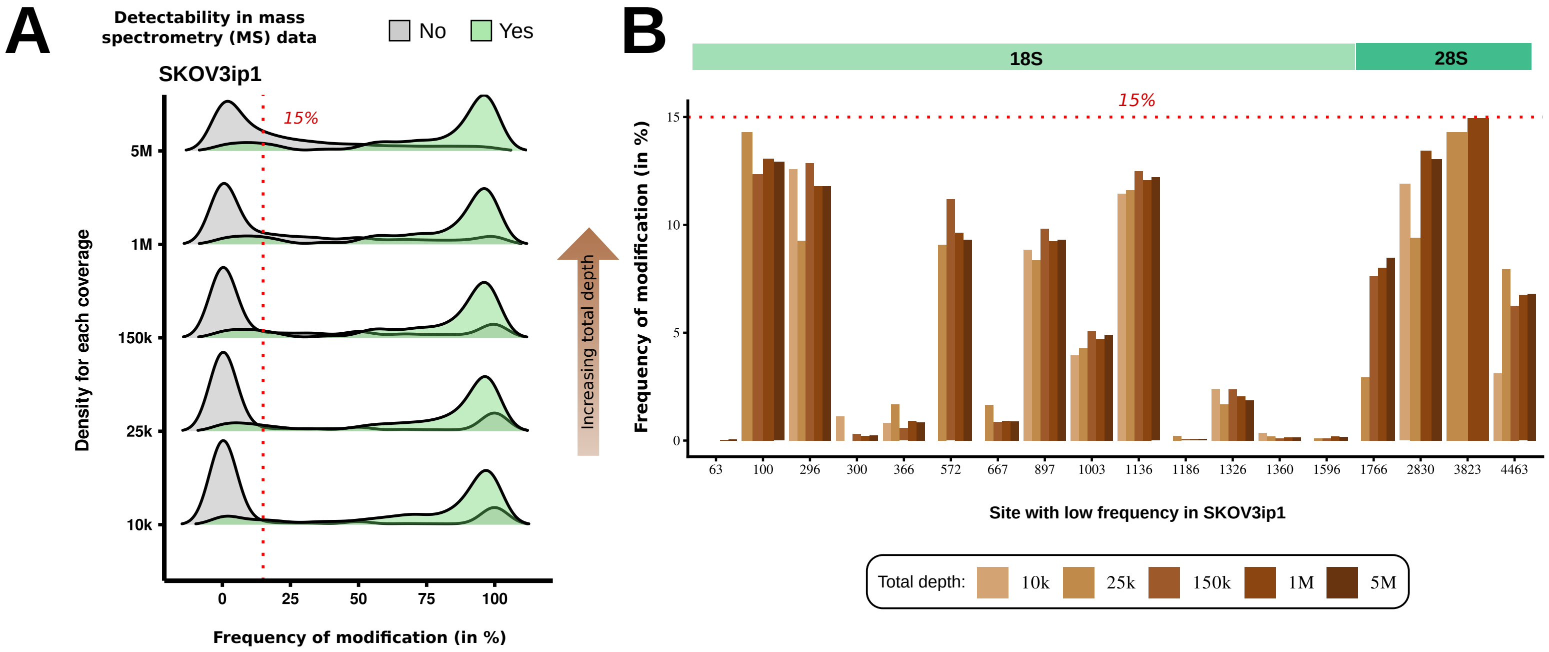


**Supplementary Figure 1. Increased read depth in SKOV3ip1 cells does not recover low frequency nanopore Ψ predictions.** (A) Kernel density plots of pseudouridine (Ψ) modification frequencies predicted from SKOV3ip1 rRNA at subsampled depths ranging from 10,000 to 5 million reads. Green shading indicates MS-validated Ψ sites; grey shading denotes sites without MS support. The red dashed line marks the 15% frequency threshold used in this study to define reliable nanopore detection. (B) Barplots showing MS-validated Ψ sites that consistently fall below the 15% nanopore detection threshold (red dashed line). Sites are ordered by genomic position and grouped by rRNA subunit, with green tiles at the top indicating their 18S or 28S location. For each site, predicted modification levels are shown across all subsampled depths (10k–5M reads; brown gradient). Despite increasing sequencing depth, these MS-validated sites remain quantified at low frequencies by nanopore sequencing.

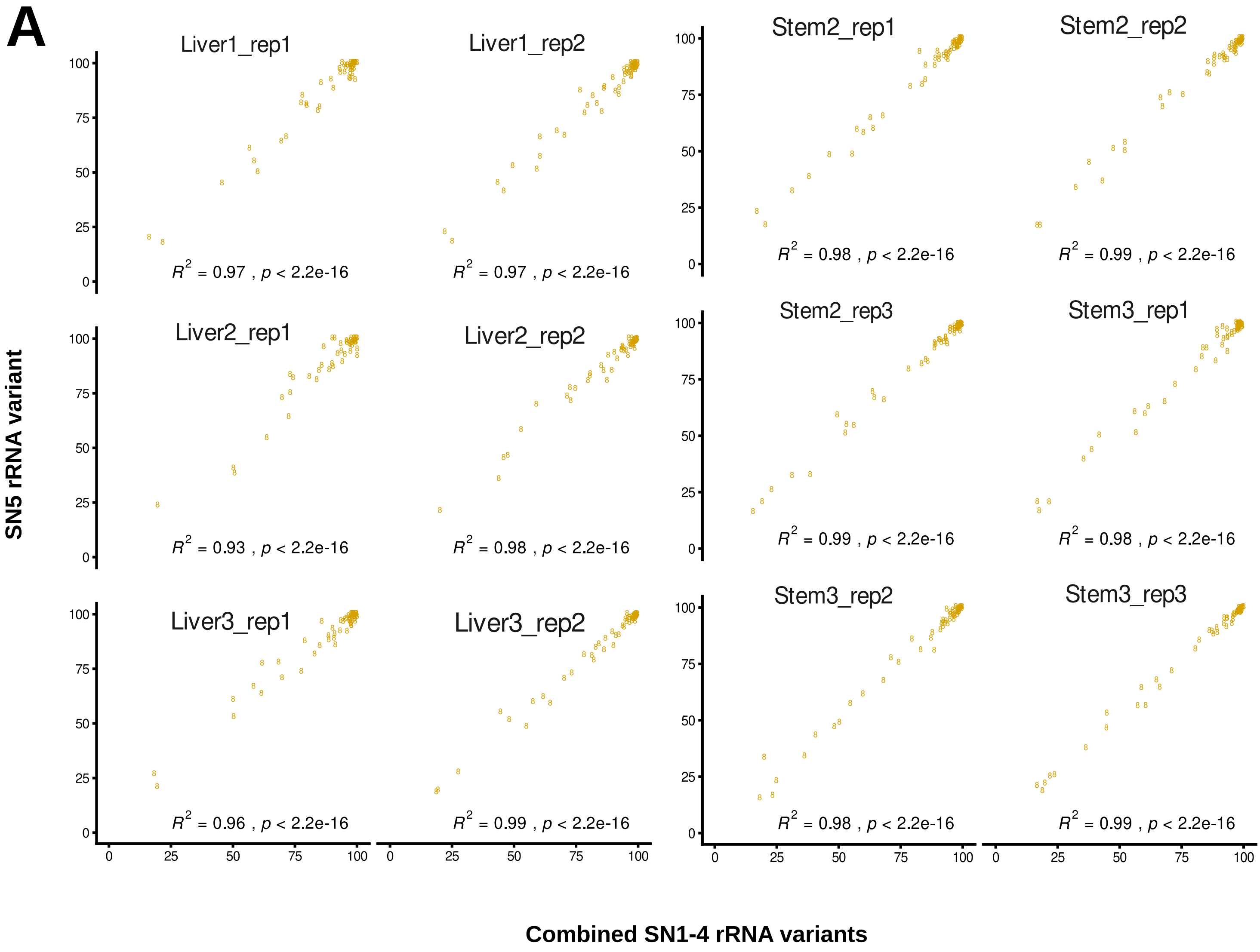


**Supplementary Figure 2. Ribosomal RNA variants of the 28S subunit have minimal impact on nanopore-based pseudouridine quantification.** Correlation plots comparing Ψ modification frequencies predicted for the 28S SN5 rRNA variant with those obtained from a weighted composite of the other 28S variants (SN1–SN4) across liver and stem cell samples. For each replicate, modification frequencies were recalculated by integrating read coverage and predicted Ψ levels from all expressed 28S variants. Each scatter plot represents one biological replicate, and the high coefficients of determination ($R^2$) indicate that variant-specific sequence differences introduce negligible variation in nanopore-based Ψ quantification.

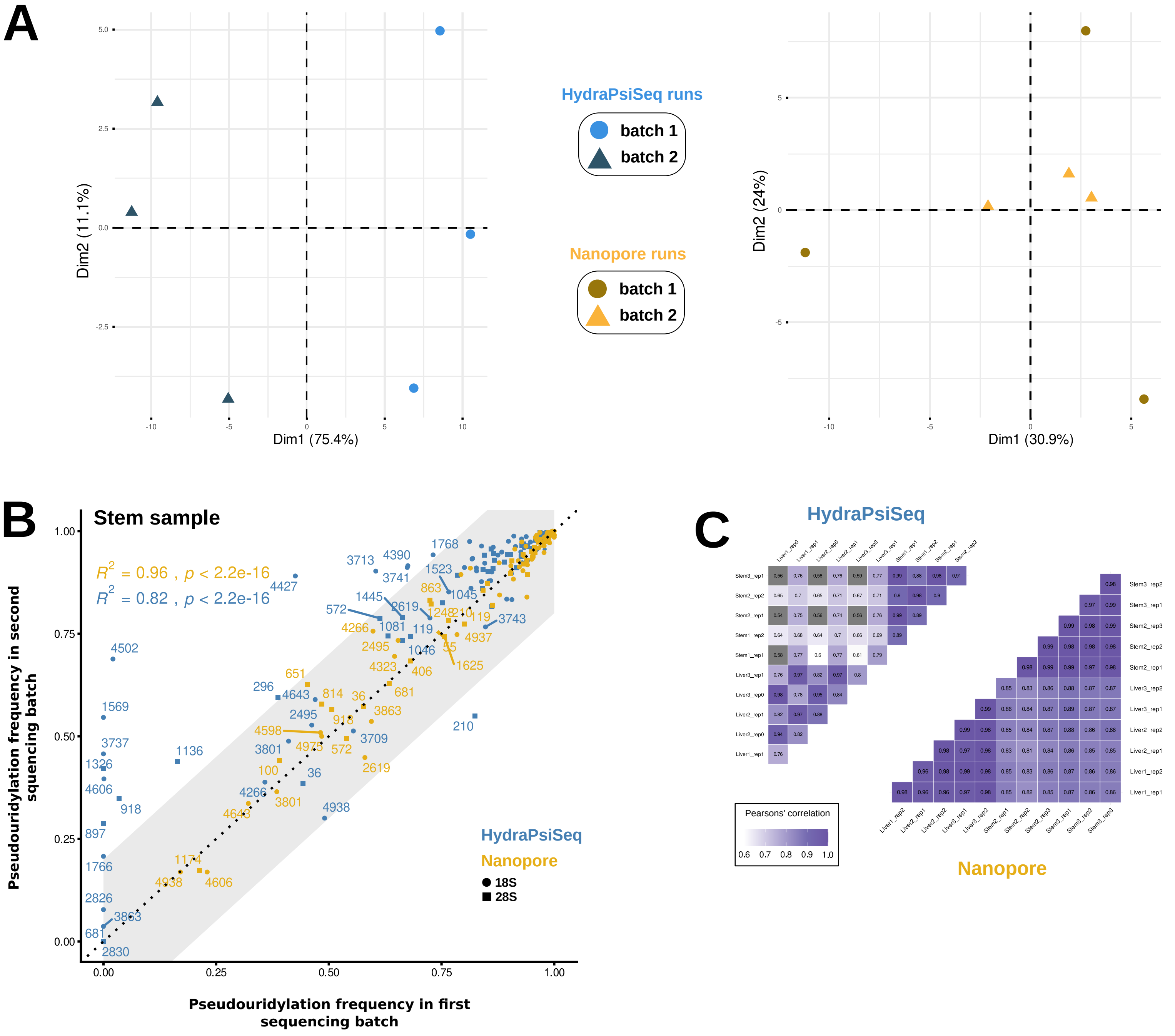


**Supplementary Figure 3. Cross-batch and cross-method reproducibility highlights higher stability of nanopore quantification.** (A) Principal component analysis of liver replicates for HydraPsiSeq (left) and nanopore (right). Points represent individual replicates, colored by method, and shaped by sequencing batches. Percent variance explained by each dimension is indicated in parentheses. HydraPsiSeq replicates show pronounced batch-driven separation along Dim1, whereas nanopore replicates cluster tightly regardless of batch. (B) Scatter plots comparing pseudouridine frequencies measured in two independent sequencing batches for the same hIPSC sample, shown separately for nanopore and HydraPsiSeq. Each point represents an rRNA site, labeled by its genomic position and colored by method (yellow for nanopore; blue for HydraPsiSeq). The grey band indicates a ±20% deviation from the identity line (x = y). Correlation coefficients ($R^2$) and p-values for each method are displayed in the upper-left corner. Nanopore estimates cluster tightly within the deviation band, whereas HydraPsiSeq measurements show substantial dispersion, particularly at low-frequency sites. (C) Pairwise Pearson correlation matrices for all liver and hIPSC replicates quantified by HydraPsiSeq (upper left matrix) and nanopore (lower right matrix). Each tile displays the correlation between two samples. Nanopore measurements show uniformly high between-sample correlations, while HydraPsiSeq exhibits greater variability across replicates and batches.

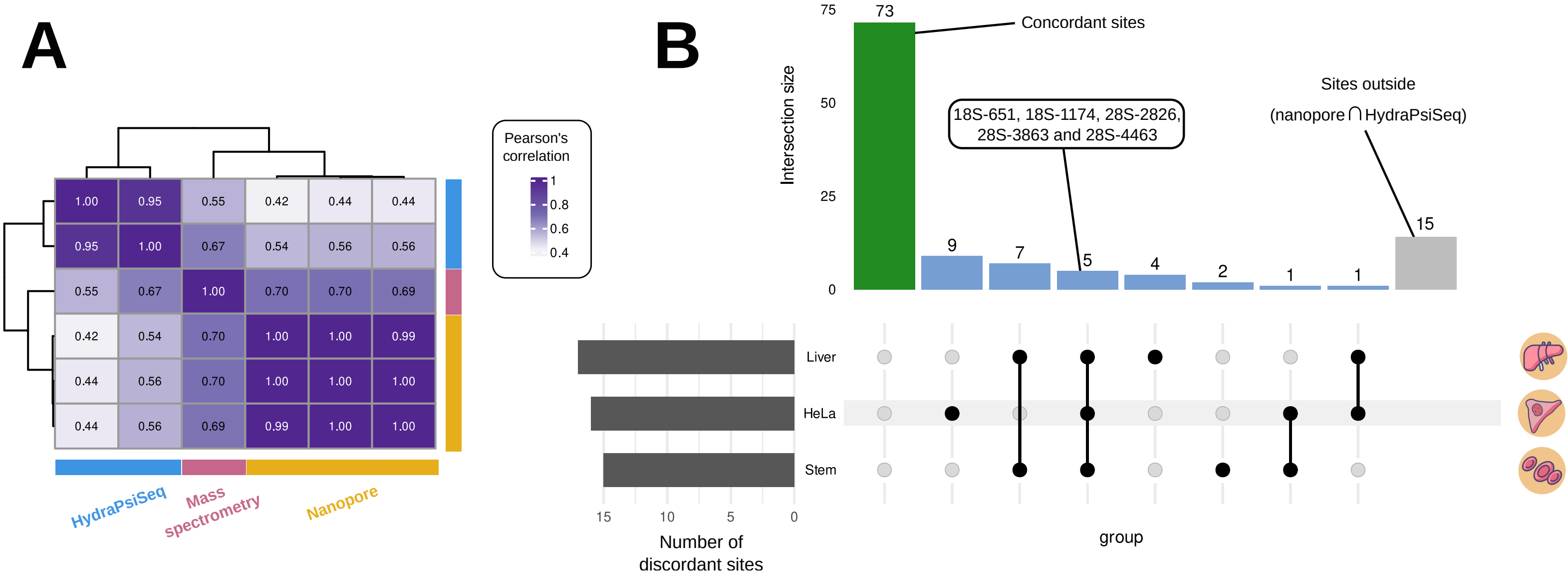


**Supplementary Figure 4. Agreements for pseudouridine frequencies in HeLa and summary points of contention across biological contexts.** (A) Correlation matrix built on pseudouridine fractions predicted by mass spectrometry, HydraPsiSeq and nanopore (respectively rectangles in pink, blue and yellow). Pearson's correlation coefficient is written on each tile of the heatmap. (B) Upset plot summarizing disagreements between HydraPsiSeq and nanopore on their quantification of pseudouridines in rRNA. On the top, bars are proportional to the number of sites concerned and colored in green if both methods agree, in pale blue if they disagree and in grey if the sites are outside their range of detection. For the triple intersection, a list of problematic sites is specified.

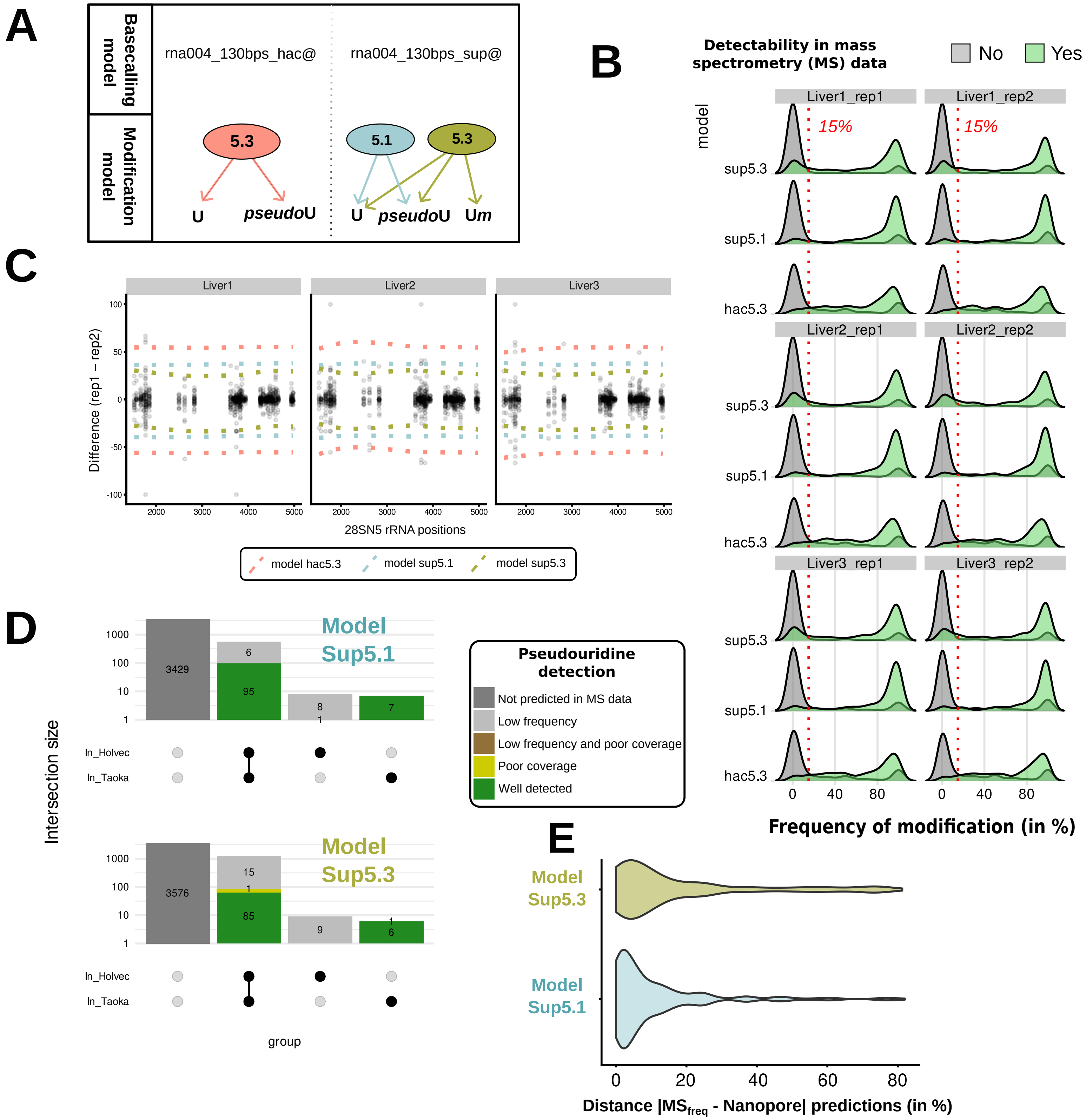


**Supplementary Figure 5. Comparison of modification-aware models sup and hac version 5.1 and 5.3 for nanopore-based pseudouridine quantification.** (A) Overview of the pseudouridine compatible dorado rna004_130bps basecalling workflows, which include two modification-aware models: model 5.1 (gold), trained to distinguish Ψ from U; and model 5.3 with two modes: (i) hac (in brown) also calling Ψ from U and (ii) sup trained to detect Ψ, 2′-O-methyl-U, and U. (B) Kernel density distributions of predicted modification frequencies for liver replicates using each model. For each site, green shading indicates positions confirmed by mass spectrometry, whereas grey denotes sites without MS support. The red dashed line marks the 15% modification frequency threshold used in this study. (C) Bland–Altman plots showing per-site differences between technical replicates (rep1–rep2) across three liver samples for rRNA variant 28SN5. Dotted lines represent the 95% confidence interval. Models sup 5.1 and 5.3 exhibit narrower limits of agreement, indicating close reproducibility relative to model hac 5.3. (D) Classification of (Ψ) sites predicted by model sup5.1 (top) or sup5.3 (bottom). Intersections per dataset are summarized with the barplot above for each model. (E) Violin plots comparing the distribution of site-wise absolute distances between nanopore models sup5.1 and sup5.3 predictions against the average of mass spectrometry fractions in HeLa cells.
